# Inter-leaflet Organization of Membrane Nanodomains: What Can(not) Be Resolved by FRET?

**DOI:** 10.1101/2022.10.31.514022

**Authors:** Barbora Chmelová, David Davidović, Radek Šachl

## Abstract

Plasma membranes as well as their simplified model systems show an inherent nanoscale heterogeneity. As a result of strong interleaflet interactions, these nanoheterogeneities (called here lipid nanodomains) can be found in perfect registration (*i.e.* nanodomains in the inner leaflet are registered with the nanodomains in the outer leaflet). Alternatively, they might be inter-leaflet independent, anti-registered or located asymmetrically in one bilayer leaflet only. To distinguish these scenarios from each other appears to be an experimental challenge. In this work, we analyzed the potential of Förster resonance energy transfer (FRET) to characterize inter-leaflet organization of nanodomains. We generated *in-silico* time-resolved fluorescence decays for a large set of virtual as well as real donor/acceptor pairs distributed over the bilayer containing registered, independent, anti-registered or asymmetrically distributed nanodomains. In this way, we were able to identify conditions that gave satisfactory or unsatisfactory resolution. Overall, FRET appears as a robust method that - when using D/A pairs with good characteristics - yields otherwise difficult-to-reach characteristics of membrane lipid nanodomains.

**STATEMENT OF SIGNIFICANCE:** This work first explores the potential of Förster resonance energy transfer (FRET) to characterize inter-leaflet nanodomain coupling and then shows how a FRET experiment can designed to achieve optimal resolution towards nanodomain coupling. Importantly, the analysis identifies as the most critical the following parameters fundamentally affecting the resolution of FRET: the Förster radius and its value related to the inter-layer distance at which donors and acceptors in the opposing membrane leaflets are separated from each other and the donor and acceptor partition coefficients characterizing their distribution between the domain and nondomain region. By setting these parameters correctly, FRET allows for the characterization of inter-leaflet nanodomain organization with unprecedented detail.

## INTRODUCTION

The ongoing intensive research of the organisation of plasma membranes suggests that they are nanoscopically heterogeneous both in their structure and chemical composition.(1–5) Up to now, the formation of membrane heterogeneities, known in literature as lipid nanodomains, has been observed not only in cellular membranes but also in their synthetic models, comprising supported phospholipid bilayers (SPBs), free-standing membranes of giant unilamellar vesicles (GUVs) or other (multi)-lamellar structures.(2, 6, 7) These nanodomains exist even in lipid mixtures containing solely two distinct types of lipids.(8) Although the nanodomains have been characterized so far by diverse experimental approaches(9), their features and importance for membrane related biological functions are still mostly unknown.

Considering that plasma membranes consist of two lipid layers that are in close contact, it is quite likely that the nanodomains in one layer will affect the positions of the nanodomains in the other layer. In principle, the following hypothetical scenarios may arise (**Figure 1**); 1) Nanodomains are perfectly registered across the bilayer leaflets, meaning that the nanodomains in the inner leaflet occupy the same lateral positions as the nanodomains in the outer leaflet **(Figure 1 A**); 2) the nanodomains exist in both leaflets and are fully independent of each other **(Figure 1 B**), 3) nanodomains are anti-registered **(Figure 1 C**), implying that the nanodomains in both leaflets avoid each other, thus, the nanodomains in the inner leaflet cannot occupy the lateral positions taken by the nanodomains in the outer leaflet, and vice versa. And 4) nanodomains are formed in an asymmetric manner in one leaflet only **(Figure 1 D**). Due to the low thickness of the lipid bilayer of only a few nanometers and small nanodomain size that is close to or below the resolution of optical microscopes, it is experimentally challenging to distinguish these possible scenarios from each other and thus to find out how nanodomains are organized between both bilayer leaflets.

**Figure 1:**
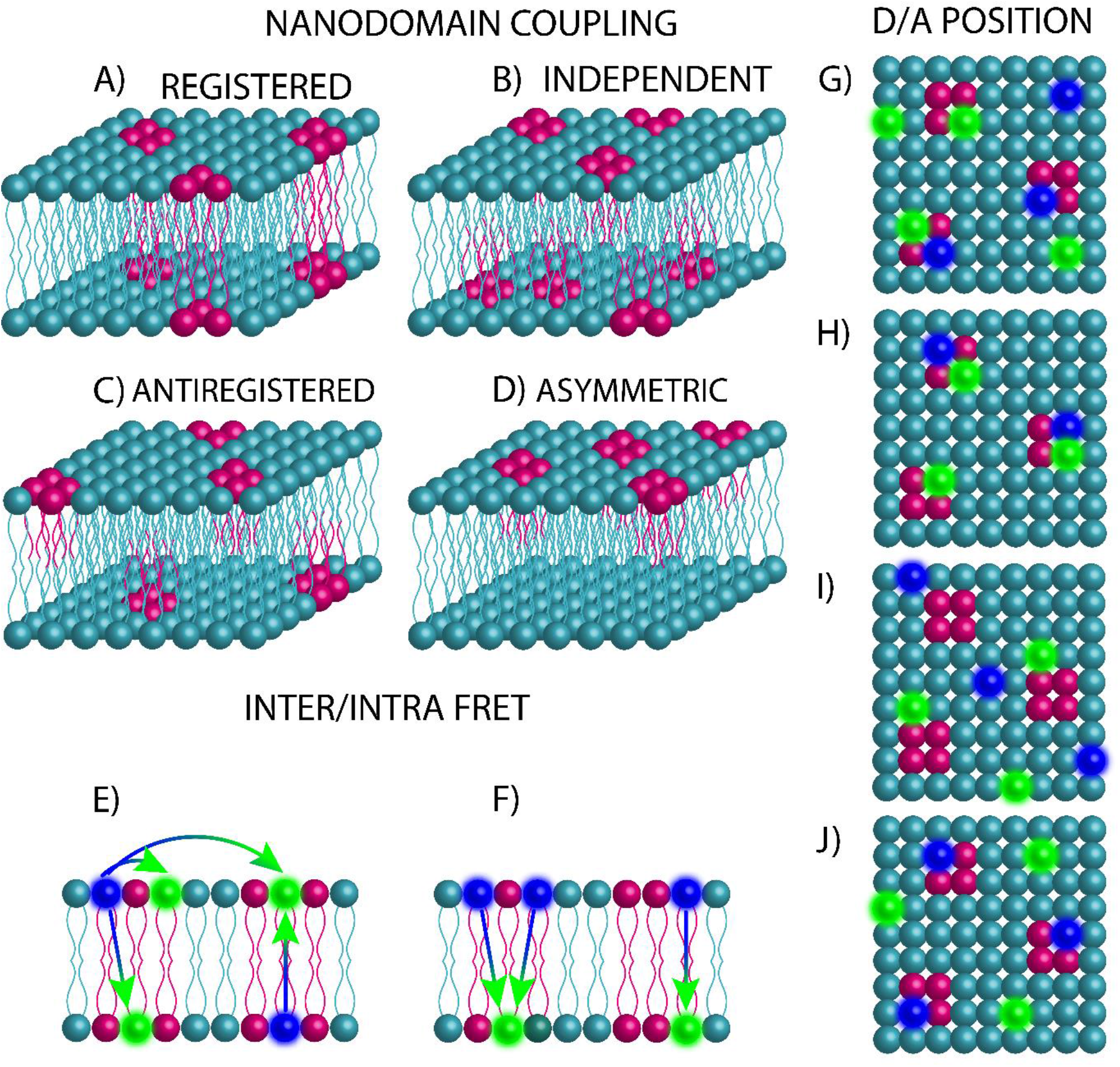
(A-D) Inter-leaflet organisation of lipid nanodomains (depicted in pink): (A) registered (= inter-leaflet coupled), (B) inter-leaflet independent, (C) anti-registered nanodomains or (D) nanodomains formed asymmetrically in one bilayer leaflet. (E-F): Distribution of D (blue spheres) and A (green spheres) in respect to both bilayer leaflets: (E) symmetric where both *intra*- and *inter*-FRET processes occur and (F) asymmetric where D and A occupy opposite leaflets. Under such circumstances, *intra*-FRET is eliminated, and only *inter*-FRET can take place. (G-J) Distribution of D and A relatively to the nanodomains: (G) both D and A have equal affinity to nanodomains and the region outside of them corresponding to ‘Case 0’, (H) both D and A have pronounced affinity to nanodomains (*K*_D_(D/A) > 1, Case I), (I) both D and A have low affinity to nanodomains (*K*_D_(D/A) < 1, Case II) and (J) D have increased affinity to nanodomains (*K*_D_(D) > 1) whereas A are excluded from them (*K*_D_(A) < 1, Case III).

**Figure 2:**
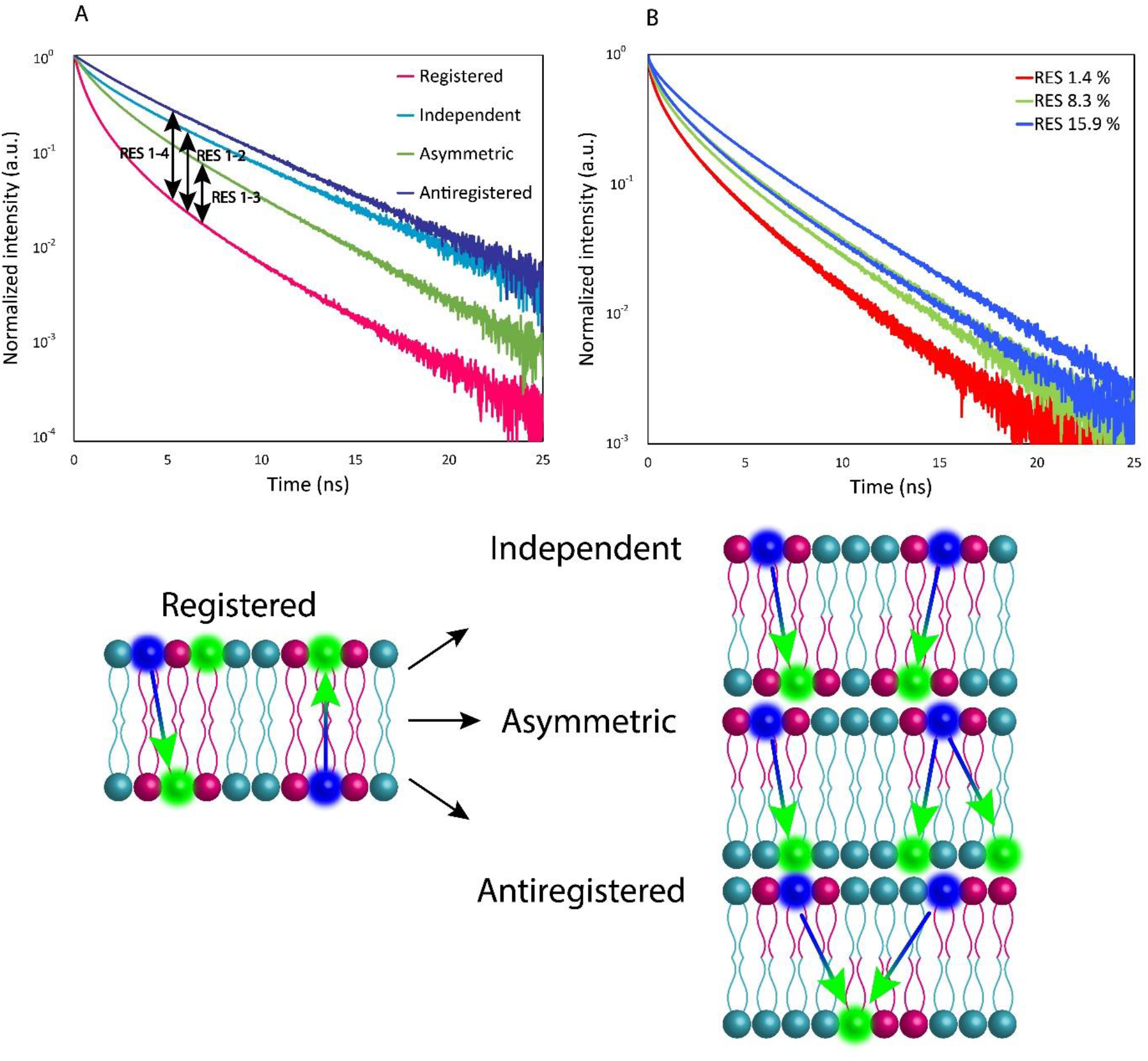
The meaning of RES parameter. In the case outlined on panel A, the parameter expresses the difference between the fluorescence decay generated for the system containing registered nanodomains (solid magenta curve) and the systems with either independent (dotted cyan curve, yielding the value RES (1–2)), anti-registered (dashed blue curve, yielding the value RES (1–3)) or asymmetrically distributed nanodomains (dash-dotted green curve, yielding the value RES (1–4)). On panel B, time-resolved fluorescence decays corresponding to RES = 1.14% (red decays), RES = 8.3% (green decays) and RES = 15.9% (blue decays) are displayed.

Recently, we introduced a new fluorescence spectroscopy method MC-FRET (Förster Resonance Energy Transfer analysed by Monte-Carlo simulations) for the characterization of inter-leaflet organization of lipid nanodomains and documented on a few specific cases that the nanodomains of variable sizes between 10 and 160 nm are perfectly registered across both leaflets.(10–12) According to our knowledge, MC-FRET is the only up-to-date available experimental technique that can resolve inter-leaflet coupled from independent nanodomains in free-standing model lipid bilayers. Its applicability to the plasma membranes of living cells will need to be tested in future.

The resolution of this method hangs on several parameters.(8, 13, 14) In particular, it depends on the intrinsic properties of the chosen donor (D)-acceptor (A) pair, including the affinity of D and A to the nanodomains (characterized by their partition coefficient *K*_D_(D/A)), their Förster radius, the distance between the D and A planes, or the distribution of D and A between both leaflets (**Figure 1**). Thus, not every D/A pair is suitable for the characterization of nanodomain coupling, and each of them has specific limits in what it can(not) resolve.

In this work, we employed Monte-Carlo (MC) simulations and generated time-resolved fluorescence decays for a large ensemble of virtual D/A pairs having various properties (see the paragraph above) as well as real D/A pairs that have already turned out to be useful in the characterization of membrane nanodomains. All of these pairs were distributed on the bilayers containing registered, independent, anti-registered or asymmetrically distributed nanodomains. In this way, we could investigate the limits of FRET in the characterization of nanodomain coupling.

We show that despite the limited choice of suitable D/A pairs and their generally low affinity to the (non)domain region which decreases the resolution, the probe partition coefficients are high/low enough to allow for characterization of nanodomain organization in a broad range of nanodomain radii *R/R*_0_ ≥ 2 (*R*_0_ denotes the Förster radius) and relative surface area occupied by nanodomains *Area* ≥ 10 % with unprecedented detail. Care should be taken when choosing D/A pairs with a low Förster radius or when resolving independent from anti-registered nanodomains, since low FRET resolution is expected in these cases. Importantly, the analysis identifies the relatively popular D/A pairs consisting of NBD-DPPE (D) and Rhodamine-DOPE (A) or Bodipy-FL-GM1 (D) and Bodipy-564/570-GM1 (A) as the most efficient ones.

## METHODOLOGY

### Generation of time-resolved fluorescence decays by Monte-Carlo simulations

Monte-Carlo simulations were used in this work to simulate FRET in nanoscopically heterogeneous bilayers separated into two distinct regions: a domain and a nondomain region. By intention, we make no assumptions about the properties or composition of the domain and non-domain regions. The results of the simulations can thus be applied both to the case where the domains are more ordered than the surroundings (having for instance a liquid ordered or gel character), or, conversely, to the situation where the surroundings are more ordered. In the simulations, the nanodomains were assumed to be circular in shape and uniform in size with the nanodomain radius 〈*R*〉. The nanodomains in both bilayer leaflets were generated to be A) registered, B) independent, C) anti-registered or D) nanodomains were localized in one leaflet only and D and A were distributed either symmetrically in both leaflets or asymmetrically in opposite bilayer leaflets (**Figure 1**). The simulations were performed in the simulation box with dimensions of 10*R*_0_ × 10*R*_0_ and its 8 copies mimicking periodic boundary conditions. The entire process is initiated with generation of a defined number of nanodomains corresponding to the surface density 〈*Area*〉 on the bilayer surface. In the next step, D and A are distributed between nanodomains and the remaining bilayer part according to their partition coefficient *K*_D_(D) and *K*_D_(A) at the D/A to lipid ratio of 1:200. Of note, the dependence of the FRET resolution on the A to lipid ratio is fairly flat between 1:200 and 1:1000, thus, any value selected within this range guarantees a good FRET resolution (see **Figure S3**). *R*_0_ is assumed to be constant in the domain and nondomain region. The *K*_D_ is defined as:

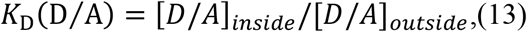

where [D/A_inside_] and [D/A_outside_] denote donor/acceptor surface concentrations inside or outside of the nanodomains, respectively.(13) Whereas D are located in the central box only at the probe to lipid ratio 1:200, A are placed into all 9 boxes at the same probe to lipid ratio to mimic the bilayers that are infinitely large. Then, a donor is randomly excited and the time at which energy transfer takes place calculated. The overall energy transfer rate Ω_i_ modulates the process according to Δ*t_i_* = –*lnγ/Ω*_i_, where *γ* is a randomly generated number between 0-1. The outcome of each simulation step is the time interval Δ*t_i_* between the excitation and energy transfer event. By constructing a histogram of Δ*t_i_* intervals, the total survival probability function *G*(*t*) is obtained, and the simulated decay of D quenched by the A, *F*_DA_(*t*), is calculated: *F*_DA_(*t*) = *G*(*t*)*F*_D_(*t*). Here, *F*_D_(*t*) denotes the experimentally recorded donor decay in the absence of A which may not necessarily be monoexponential.(10, 15) Unless otherwise stated, a biexponential decay of Bodipy-FL attached to the headgroup of ganglioside GM1 was used as *F*_D_(*t*) that enters the simulations.

## RESULTS AND DISCUSSION: ANALYSING THE RESOLUTION OF FRET

MC-FRET has been developed to fit experimental time-resolved fluorescence decays of the donors quenched by acceptors with the decays generated by MC simulations.(8, 16) As we have shown by simulations and experiments,(8, 10) if fluorescent probes having increased or decreased affinity to lipid nanodomains are used in an MC-FRET experiment, shape of recorded fluorescence decays will change depending on the size of nanodomains (characterized by their average nanodomain radius 〈*R*〉), their surface density 〈*Area*〉 and, importantly, on the organization of nanodomains between both bilayer leaflets (**Figure 3**). Consequently, varying the input simulation parameters, 〈*R*〉 and 〈*Area*〉 can be determined.(10, 17)

**Figure 3:**
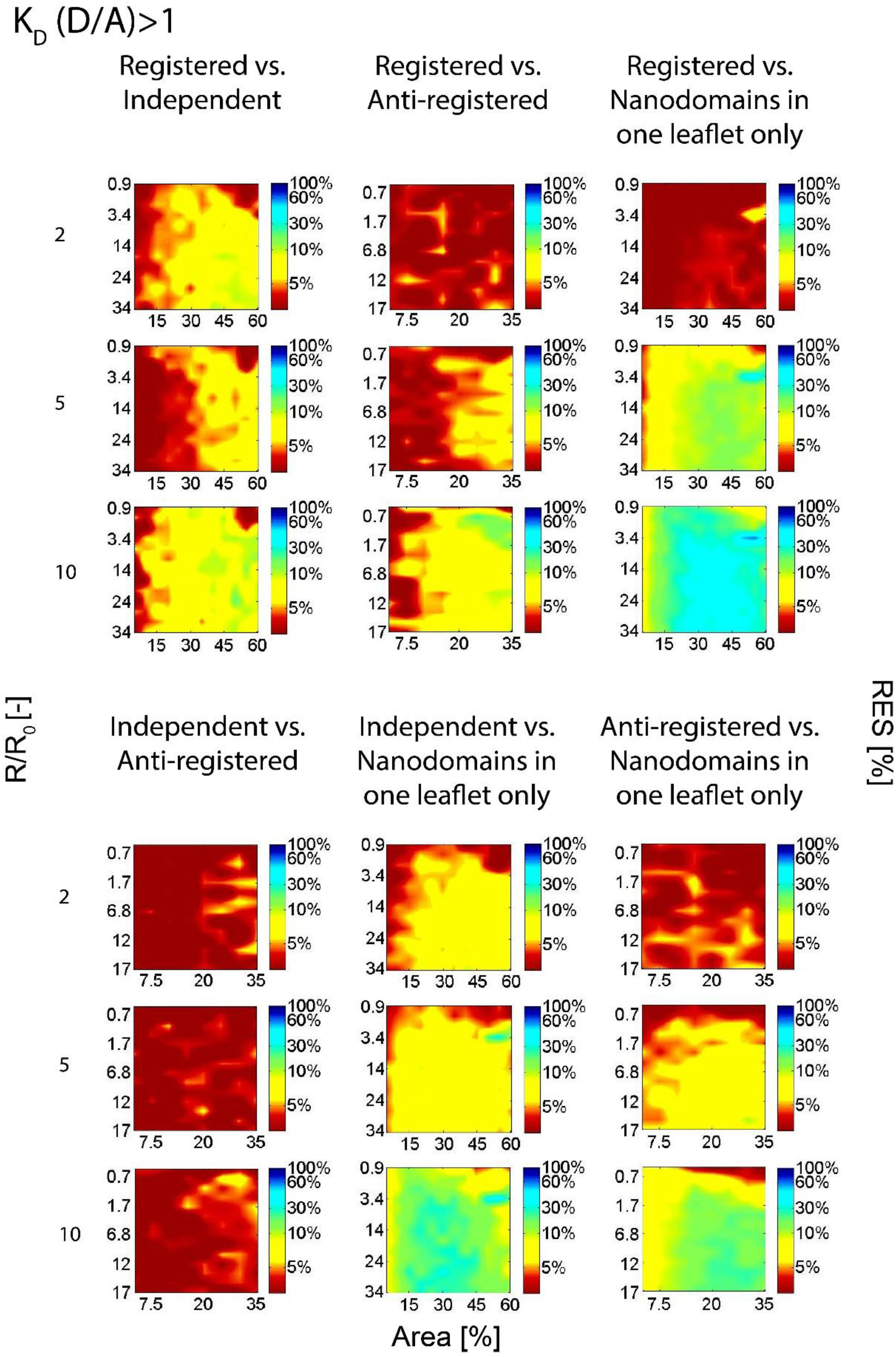
Resolution (RES) diagrams displaying the dependence of the RES parameter on the nanodomain radius (*R*) and relative surface area (*Area*) shown for the case when both D and A have increased affinity to the nanodomains (*K*_D_ (*D/A*) > 1).

Moreover, thanks to the energy transfer that occurs from one leaflet to the other one, MC-FRET offers excellent axial resolution and can be used to discriminate between the following most likely scenarios that can occur in the membrane: 1) registered **(Figure 1 A**), 2) inter-leaflet independent **(Figure 1 B**) and 3) anti-registered **(Figure 1 C**) nanodomains and 4) nanodomains formed asymmetrically in one leaflet only **(Figure 1 D**). The most probable scenario is identified by comparing the fits by means of *χ*^2^ values for scenarios 1-4.(10)

### The resolution of FRET can be characterized by ‘RES parameter’

In this work, we introduce a parameter RES, defined as

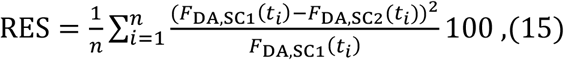

which turns out to be helpful in the characterization FRET resolution towards nanodomain coupling. RES expresses the difference between the simulated fluorescence decay for registered/independent/anti-registered or asymmetrically distributed nanodomains, *F*_DA,SC1_(*t*), and the simulated decay for one of the three alternative scenarios (see **Figure 1** for a more thorough explanation of the RES parameter); furthermore, *n* corresponds to the number of channels in the experimental decay. Thus, RES reports on the potential of (MC)-FRET to distinguish one scenario from another (for instance scenario 1 accounting for registered nanodomains from scenario 2 accounting for independent nanodomains). With the help of this parameter, resolution (RES) diagrams displaying the dependence of the RES parameter on the nanodomain radius 〈*R*〉 and surface density 〈*Area*〉 can be constructed and used to characterize the resolution of MC-FRET as follows: 1) RES ≤ 5 % (**Table 1** and red colour code on **Figure 3, Figure 4, Figure 5, Figure 6, Figure 7** and **Figure 8**), yielding very similar fluorescence decays for scenarios (SC) 1-4. At the same time, this parameter value corresponds to the relative change in the steady-state intensity of donors in the presence of acceptors, 〈*F*_DA,SC1_〉/〈*F*_DA,SC2_〉, and the intensity-weighted mean fluorescence lifetime, 〈*τ*_DA,SC1_〉/〈*τ*_DA,SC2_〉, of less than 5 % and 4 % respectively. Such conditions are unsatisfactory for the characterization of nanodomains by MC-FRET. 2) RES ∈ (5; 10) % (**Table 1** and yellow colour code on **Figure 3, Figure 4, Figure 5, Figure 6, Figure 7** and **Figure 8**), enabling the detection of nanodomains by MC-FRET. This parameter value is accompanied by 〈*F*_DA,SC1_〉/〈*F*_DA,SC2_〉 ∈ 5 – 13 % and 〈*τ*_DA,SC1_〉/〈*τ*_DA,SC2_〉 ∈ 4 – 10 %. 3) RES ∈ (10; 30) % (**Table 1** and green colour code on **Figure 3, Figure 4, Figure 5, Figure 6, Figure 7** and **Figure 8**), yielding clearly distinct fluorescence decays. This parameter value results in the change of 〈*F*_DA,SC1_〉/〈*F*_DA,SC2_〉 ∈ 13 – 60 % and 〈*τ*_DA,SC1_〉/〈*τ*_DA,SC2_〉 ∈ 10 — 30 %. 4) RES ∈ (30; 60) % (**Table 1** and cyan colour code on **Figure 3, Figure 4, Figure 5, Figure 6, Figure 7** and **Figure 8**), leading to changes in 〈*F*_DA,SC1_〉/〈*F*_DA,SC2_〉 ∈ 60 — 100 % and 〈*τ*_DA,SC1_〉/〈*τ*_DA,SC2_〉 ∈ 30 — 50 % .5) Finally, if RES > 60 % (**Table 1** and blue colour code on **Figure 3, Figure 4, Figure 5, Figure 6, Figure 7** and **Figure 8**), 〈*F*_DA,SC1_〉/〈*F*_DA,SC2_〉 > 100 % and 〈*τ*_DA,SC1_〉/〈*τ*_DA,SC2_〉 > 50 %.^(15)^

**Table 1:** RES parameter and its relation to the resolution of FRET, and the corresponding relative change in the steady state intensity and average fluorescence lifetime and the colour code used in **Figure 3, Figure 4, Figure 5, Figure 6, Figure 7** and **Figure 8.**

| RES | MC-FRET resolution | Change in the steady state intensity | Change in the average lifetime | Colour in the diagram |
| --- | --- | --- | --- | --- |
| $RES \leq 5\%$ | UNSATISFACTORY | 0-5 % | 0-4 % | RED |
| $RES \in (5;10)\%$ | SUFFICIENT | 5-13 % | 4-10 % | YELLOW |
| $RES \in (10;30)\%$ | SATISFACTORY | 13-60 % | 10-30 % | GREEN |
| $RES \in (30;60)\%$ | GOOD | 60-100 % | 30-50 % | CYAN |
| $RES \geq 60\%$ | EXCELLENT | >100 % | >50 % | BLUE |

**Figure 4:**
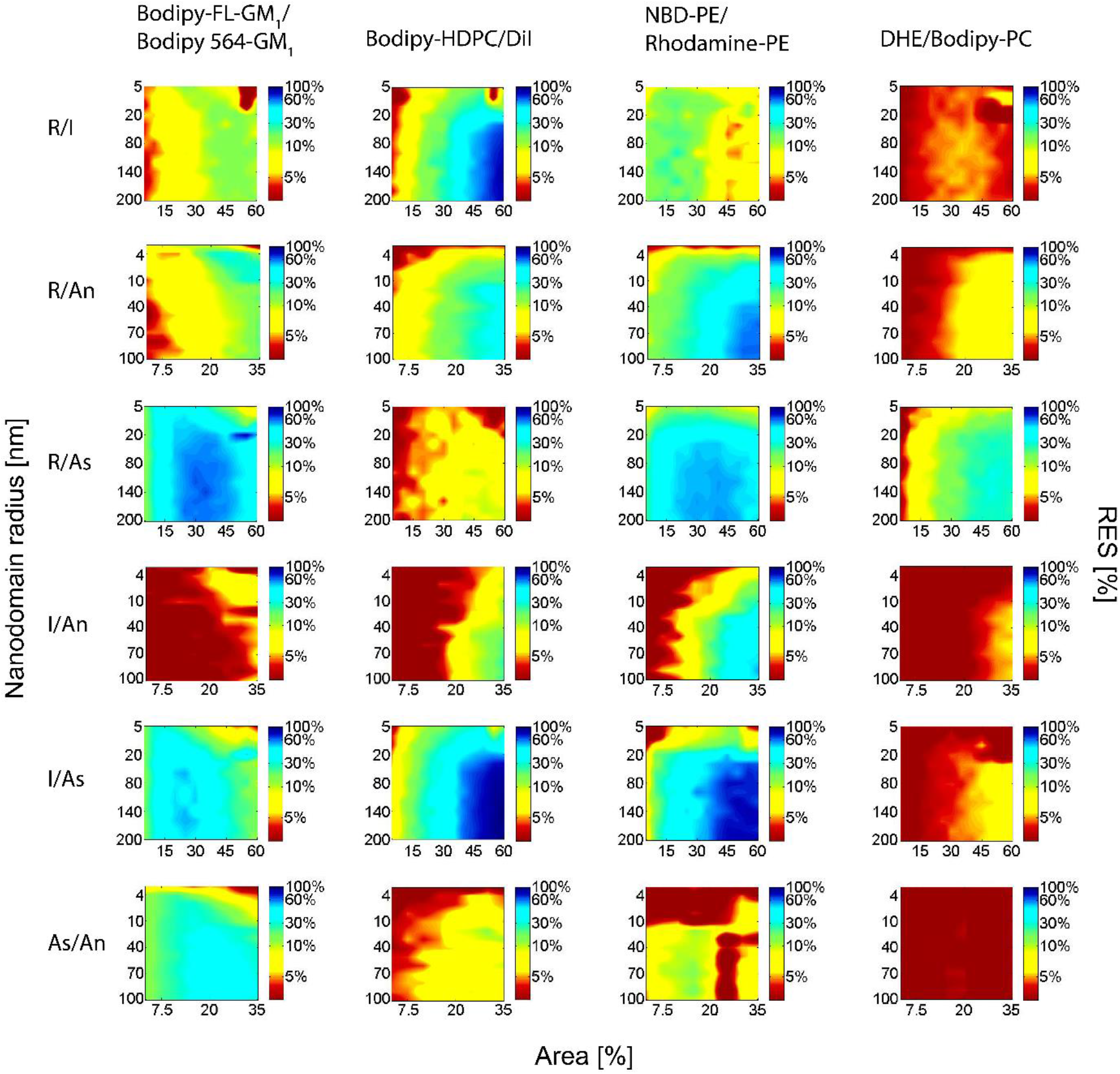
RES diagrams calculated for selected D/A pairs (see Examples I-IV in the text). D and A chromophores were located at the lipid-water interface.

**Figure 5:**
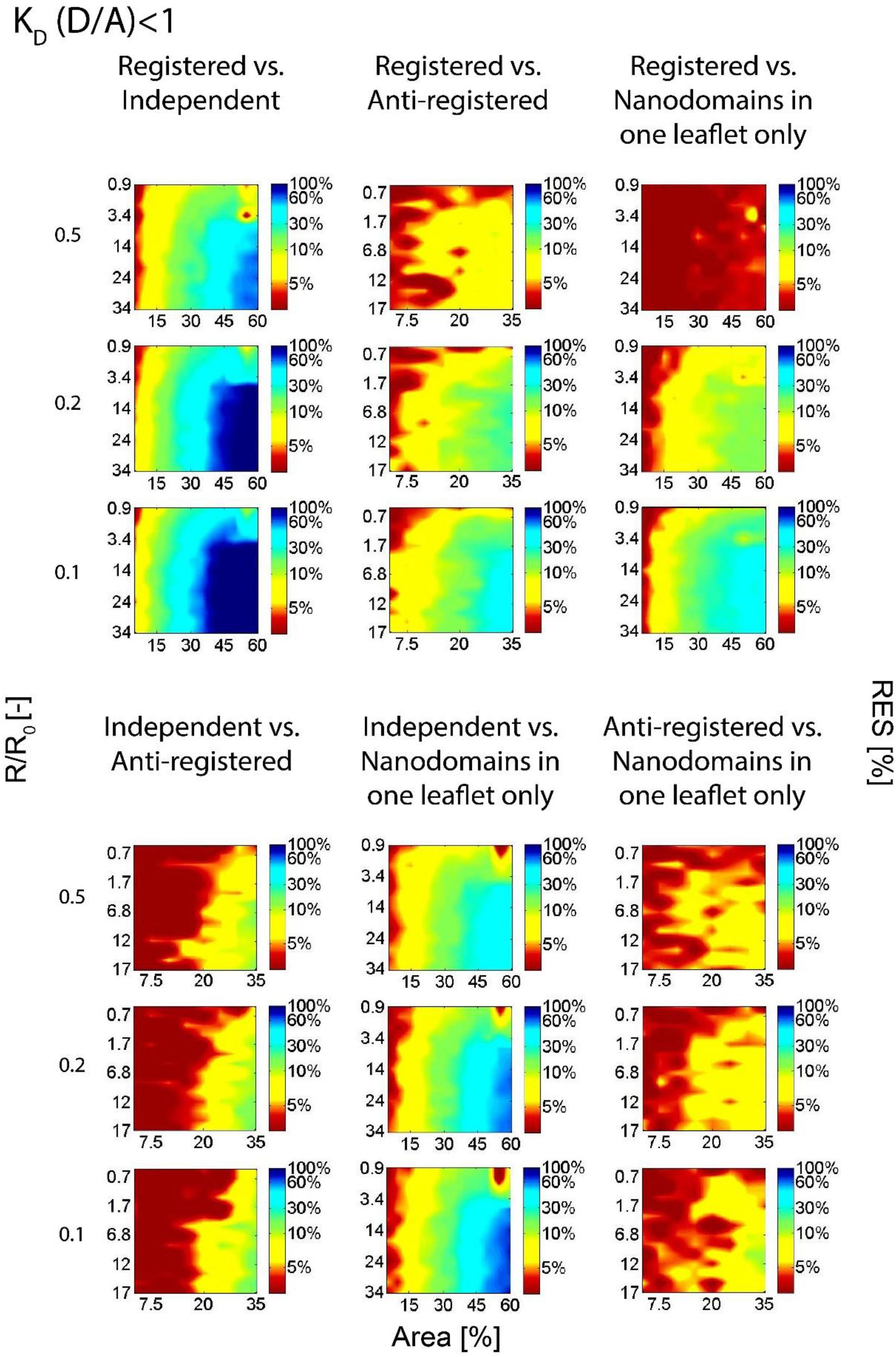
RES diagrams shown for the case when both D and A have increased affinity to the regions outside of the nanodomains (*K*_D_ (D/A) < 1).

**Figure 6:**
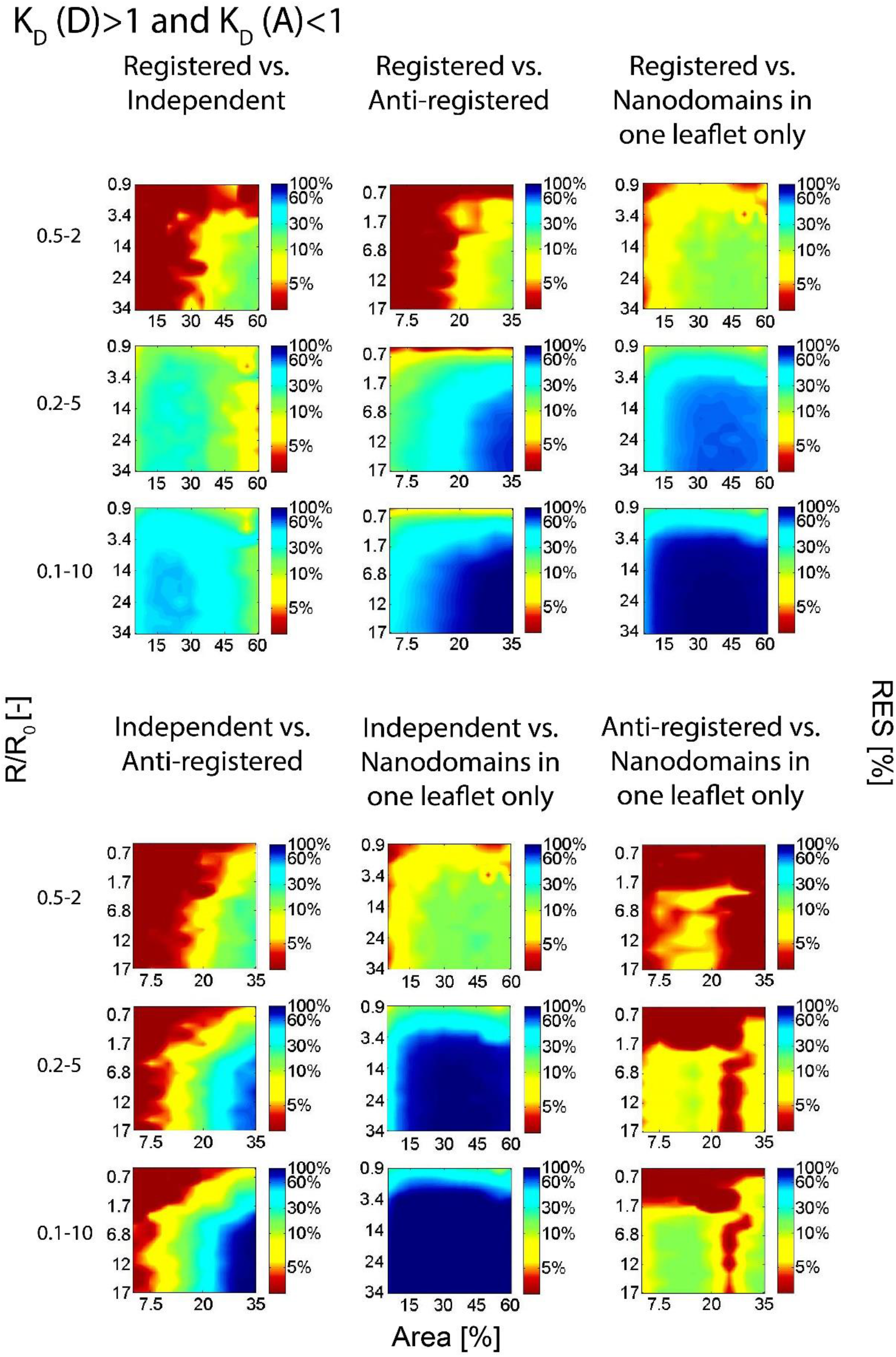
RES diagrams shown for the case when D have increased affinity to the nanodomains (*K*_D_(D) > 1) and A have increased affinity to the regions outside of the nanodomains (*K*_D_(D/A) < 1).

**Figure 7:**
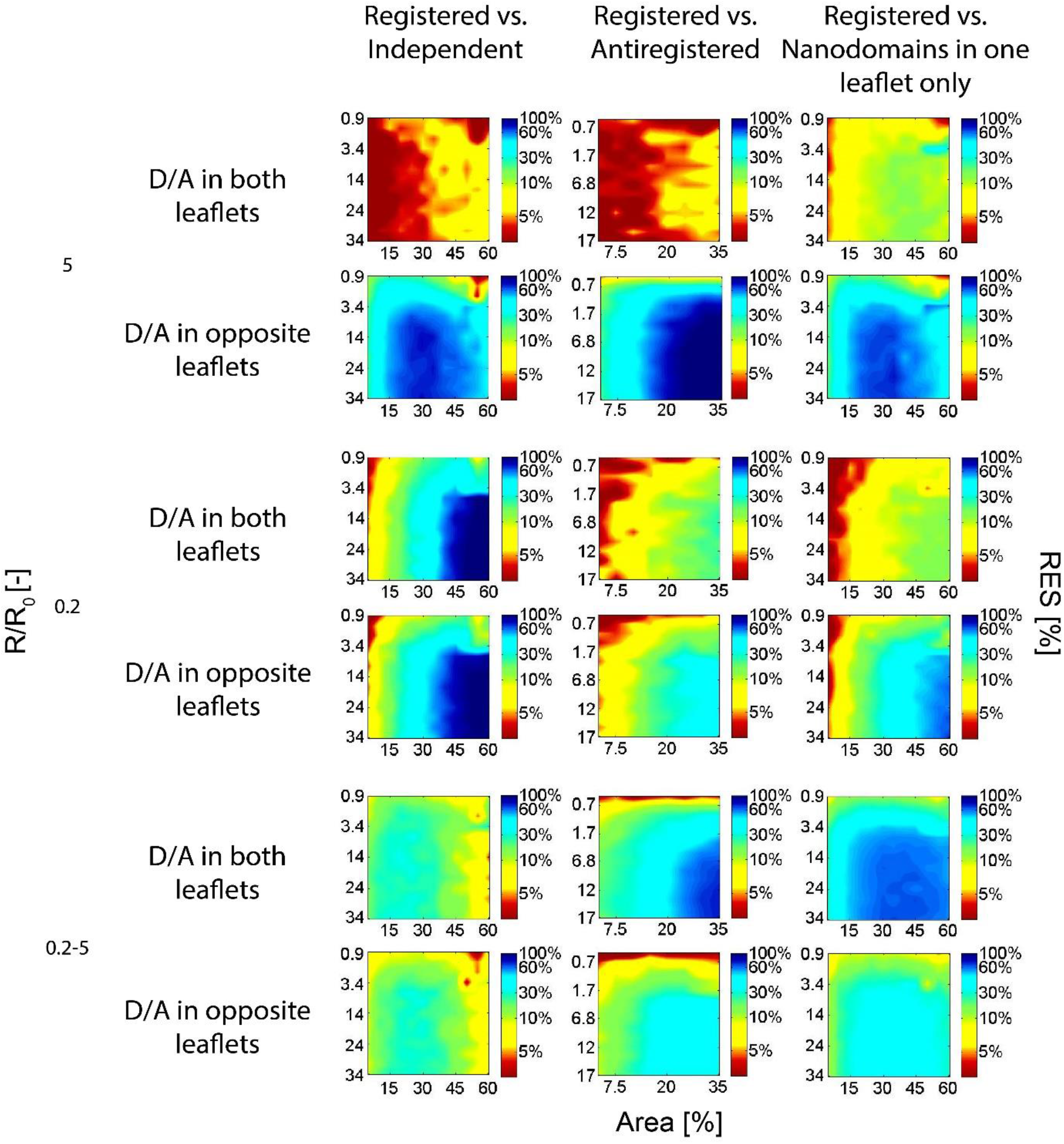
RES diagrams demonstrating the improvement of FRET resolution by distributing D and A into opposite bilayer leaflets. The simulations were performed for *K*_D_(*D/A*) = 5; *K*_D_(*D/A*) = 0.2; and *K*_D_(*D*) = 5 & *K*_D_(*A*) = 0.2. In this figure, the resolution between registered nanodomains and independent/anti-registered/asymmetrically distributed nanodomains has been investigated.

**Figure 8:**
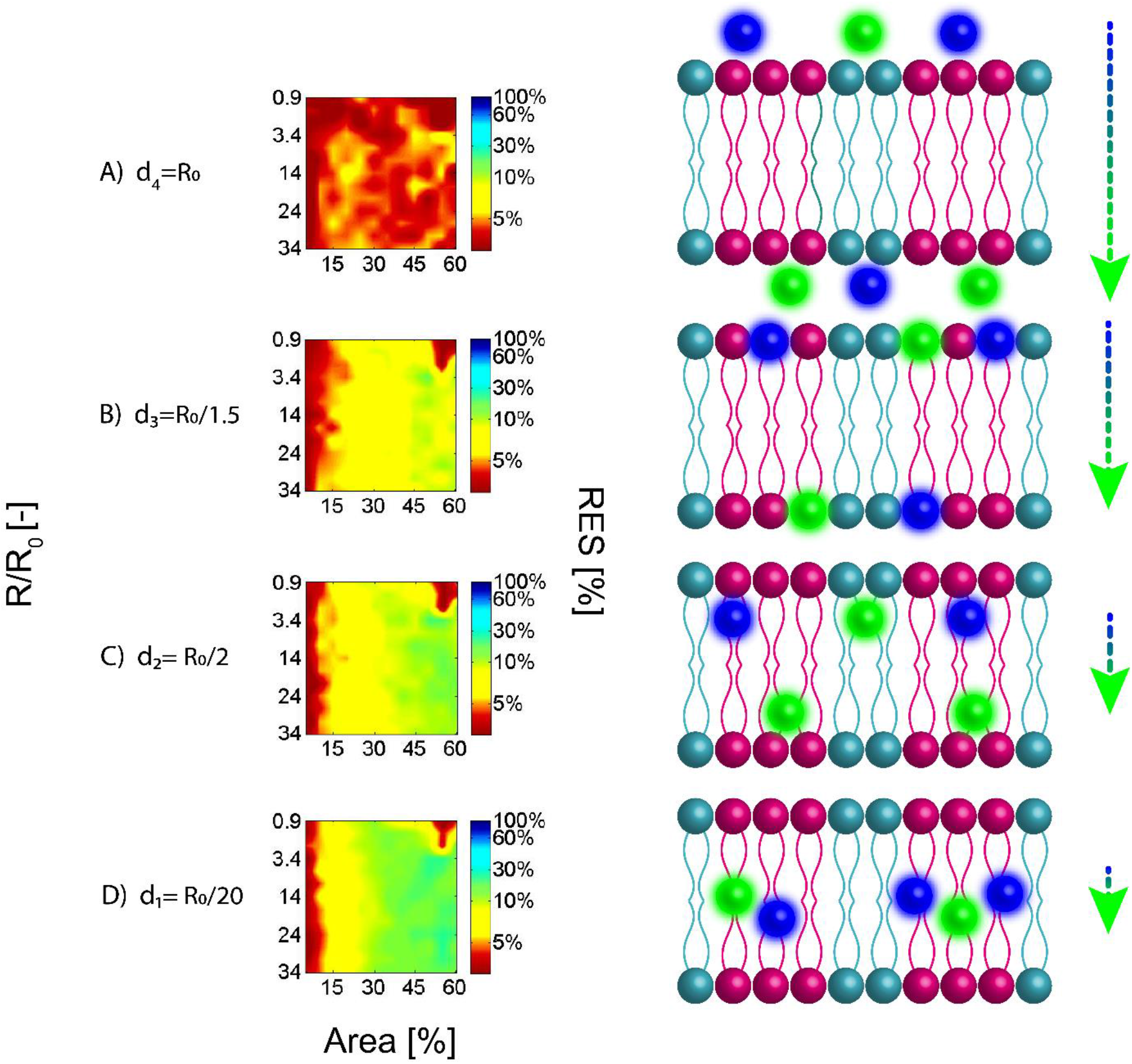
The impact of the of the inter-layer distance d on the resolution of MC-FRET. Chromophores (donors-blue and acceptors-green) with *K*_D_(D/A) = 10 were assumed to be localized along the bilayer normal in the following ways: A) fully exposed to the bulk with the inter-layer distance d = *R*_0_; B) localized close to the lipid-water interface (d = *R*_0_/1.5); C) localized below the lipid-water interface (d = *R*_0_/2) or D) deeply buried in the membrane close to the bilayer centre (d = *R*_0_/20). D and A were assumed to be localized in both bilayer leaflets; *R*_0_ = 58.7 Å **(Table 3)**

### The resolution of FRET is controlled by probes’ affinity to nanodomains

The potential of MC-FRET to characterize nanodomain coupling largely depends on which fluorescent probes are chosen as donors and acceptors of FRET, in particular it relies on the affinity of D and A to the nanodomains and the region outside of them. In this paper, this affinity is expressed in the form of partition coefficients of donors *K*_D_(D) and acceptors *K*_D_(A) (see **Methodology** for the exact definition). In principle, the following cases can arise: Case 0): *K*_D_(D) = 1 and *K*_D_(A) = 1. Consequently, D and A are distributed homogenously across the entire bilayer regardless of the presence of nanodomains **(Figure 1 G**). Such a situation allows neither for the detection of nanodomains nor characterization of their inter-leaflet arrangement using MC-FRET. Case I) *K*_D_(D) > 1 and *K*_D_(A) > 1 where both D and A are preferentially localized in nanodomains **(Figure 1 H**). Case II): *K*_D_(D)< 1 and *K*_D_(A) < 1 in which both D and A are excluded from nanodomains **(Figure 1 I**). In this and the previous case, the average distance between D and A is decreased, which results in enhanced FRET and accelerated donor relaxation kinetics as compared to Case 0. And finally, Case III): *K*_D_(D) > 1 and *K*_D_(A) < 1 or *K*_D_(D) < 1 and *K*_D_(A) > 1, which leads to accumulation of D and A in distinct bilayer regions, and, consequently, spatial separation of D from A **(Figure 1 J**). Such a probe distribution yields a lower efficiency of FRET and slower donor relaxation kinetics in comparison to Case 0. In the following text, we are going to discuss Cases I-III in more detail.

### Case I: Accumulation of D and A in nanodomains (*K*_D_(D) > 1 and *K*_D_(A) > 1) yields satisfactory resolution

In the simulations, we focused exclusively on the situations where *K*_D_(D) = *K*_D_(A) and generated data for *R*/*R*_0_ ∈ 〈0.7; 34〉, which corresponds to the nanodomain radii *R*/*R*_0_ ∈ 〈4; 200〉 nm if a typical value of *R*_0_ = 58.4 Å is used, and *Area* ∈ 〈5; 60〉%. If *R*/*R*_0_ > 34, nanodomains can be detected by classical fluorescence microscopy, which eliminates the need of using MC-FRET for nanodomain characterization. Yet, FRET could still be used in the characterization of nanodomain coupling. Moreover, if *Area* > 60%, it becomes sterically impossible to place more nanodomains into the bilayer in a way so they do not overlap. An exception is represented by anti-registered nanodomains, where it is technically impossible to achieve a density of nanodomains higher than 35% and *R*/*R*_0_ > 17. Under these conditions, more nanodomains cannot be generated in the bilayer in a way that nanodomains in both leaflets occupy distinct lateral positions.

As expected, the resolution of FRET is continuously improving as the affinity of D and A to nanodomains increases (**Figure 3**). In the main text of this paper, we only present the simulations results for *K*_D_(D, A) ∈ 〈2; 10〉, which covers the range of typical *K*_D_(D, A) values of currently available fluorescent probes (see **Table 2:** The list of fluorescent probes with either increased or decreased affinity to lipid domains. for the list of these probes). As shown in **SI**, the MC-FRET resolution continues getting better only slightly between *K*_D_(D, A) ∈ 〈10; 100〉, but beyond *K*_D_(D, A) > 100, the resolution is not significantly improved. The continuously improved resolution seems common to all six situations we examined: specifically, the situations where the potential of MC-FRET to resolve registered from independent, registered from anti-registered or registered nanodomains from the nanodomains localized in one leaflet was investigated **(Figure 3,** the upper row). Furthermore, we investigated the capacity of MC-FRET to resolve independent from anti-registered, independent nanodomains from those localized asymmetrically in one leaflet and anti-registered from asymmetrically distributed nanodomains (**Figure 3**, the lower row).

**Table 2:** The list of fluorescent probes with either increased or decreased affinity to lipid domains. Note that the *K*_D_ values reported in this table are strictly related to the listed lipid compositions and their values must be determined again for other lipid compositions.

| | Fluorophore | $K_D$ | Lipid composition | Ref |
| --- | --- | --- | --- | --- |
| Probes with affinity to the nondomain region | NBD-DLPE<sup>a</sup> | 0.21/0.41 | DLPC/DSPC<sup>b</sup> (60:40/40:60) | (19) |
|  | DiIC18(3) | 0.15/0.15 | DLPC/DSPC<sup>b</sup> (60:40/40:60) |  |
|  | BODIPY-PC (16:0) | 0.16±0.025 | bSM<sup>c</sup>/DOPC/POPC/Chol (40/40/20) | (20) |
|  | Rhodamine-DOPE | 0.37±0.06 | PSM<sup>d</sup>/POPC/Chol (33/33/33) | (21) |
|  | BODIPY-HDPC<sup>e</sup> | 0.1 | DOPC/DSPC<sup>b</sup>/Chol (35/35/30) | (22) |
|  | fast-DiI | 0.1 | DOPC/DSPC<sup>b</sup>/Chol (35/35/30) |  |
|  | DiD | 0.004±0.002 | DOPC/SM/Chol/DOPG/GM1<br>(23-68/0-45/25/5/2) | (23) |
|  | TOE<sup>f</sup> | 0.1±0.01 | bSM<sup>c</sup> /DOPC/POPC/Chol (40/40/20) | (20) |
| Probes with affinity to domain region | Bodipy FL-GM1 | 10 | DOPC/SM (85-90/10-15) | (8) |
|  | Bodipy 564/570-GM1 | 10 | DOPC/SM (85-90/10-15) |  |
|  | Bodipy FL-GM1 | ≥20 | DOPC/Chol/SM (65-70/25/5-10) |  |
|  | Bodipy 564/570-GM1 | ≥20 | DOPC/Chol/SM (65-70/25/5-10) |  |
|  | Alexa 488-CTxB<sup>g</sup> | 6±3 | DOPC/SM/Chol/DOPG/GM1<br>(23-68/0-45/25/5/2) | (23) |
|  | NBD-DPPE | 4.3±1.2 | PSM<sup>d</sup>/POPC/Chol (33/33/33) | (21) |
|  | DHE | 3.7 | bSM<sup>c</sup>/DOPC/POPC/Chol (40/40/20) | (20) |
|  | NBD-DHPE | 2.2 | DPhPC<sup>h</sup>/DPPC/Chol (40/35/25) | (24) |

<sup>a</sup> N-(7-nitrobenz-2-oxa-1,3-diazol-4-yl)-dilauroylphosphatidylethanolamine
<sup>b</sup> Distearoylphosphatidylcholine
<sup>c</sup> Sphingomyelin - brain
<sup>d</sup> Palmitoyl sphingomyelin
<sup>e</sup> 2-(4,4-difluoro-5,7-dimethyl-4-bora-3a,4a-diaza-s-indacene-3-pentanoyl)-1-hexadecanoyl-sn-glycero-3-phosphocholine
<sup>f</sup> tryptophan oleoyl ester
<sup>g</sup> Alexa Fluor 488-cholera toxin
<sup>h</sup> Diphytanoylphosphatidyl choline

It is obvious from the resolution maps that even for D and A having moderate affinity to nanodomains (*K*_D_(D/A) >5) a satisfactory resolution, *i.e.* a resolution where the red color in the resolution diagrams represents a minor component, is achieved for all analyzed situations. The only exception is the scenario where independent nanodomains are resolved from antiregistered ones. In this specific case, the nanodomains are distributed over the bilayer surface in a very similar manner, which results in similar time-resolved fluorescence decays, and consequently poor resolution of MC-FRET. As a rule, it appears most difficult to study nanodomain coupling if they occupy only a small part of the bilayer *Area* ≤ 10%, independently of the nanodomain size.

### Example I: Bodipy-FL-GM1/ Bodipy-564/570-GM_1_ D/A pair (K_D_(D/A) ≥ 20, R_0_ = 58.4 Å)

This D/A pair consisting of fluorescently labelled gangliosides GM_1_ (**Table 3** and **Figure 4**) exhibits the highest experimentally determined affininty to lipid nanodomains and has already been used in a variety of experimental studies, for instance in the detection and characterization of DOPC/Chol/SM nanodomains, ganglioside nanodomains or nanodomains contaning oxidized phospholipids (8, 10, 11, 13, 17, 18).

**Table 3:** The main properties (including the probe affinity to the (non)domain region and the Förster radius *R*_0_) of the D/A pairs presented in Examples I - IV. Note that the *K_D_* values reported in this table are strictly related to the listed lipid compositions and their values must be determined again for other lipid compositions.

| D | A | $K_D(D)$ | $K_D(A)$ | Lipid composition | $R_0$ (nm) | Ref |
| --- | --- | --- | --- | --- | --- | --- |
| DHE | Bodipy-PC<br>(16:0) | 3.7 | $0.16 \pm 0.025$ | bSM/DOPC/<br>POPC/Chol (40/40/20) | 2.8 | (20) |
| Bodipy FL -<br>GM1 | Bodipy 564 -<br>GM1 | $\geq 20$ | $\geq 20$ | DOPC/Chol/SM<br>(65-70/25/5-10) | 5.87 | (8) |
| NBD-DPPE | Rhodamine-DOPE | 4.3±1.2 | 0.37±0.06 | PSM/POPC/Chol (33:33:33) | 6.41 | (21) |
| BODIPY-HDPC | Fast DiI | 0.1 | 0.1 | DOPC/DSPC/Chol (35/35/30) | 6.5 | (22) |

In our recent work, we used this D/A pair to study interleaflet organization of nanodomains by MC-FRET.(10) More specifically, we fitted experimentally recorded decays by the models assuming registered, independent and anti-registered nanodomains and identified the following global minima in the *χ*^2^ space: *χ*^2^(REG) = 1.94 (R = 78 ± 17 nm,Areα = (63 ± 5)%); *χ*^2^(INDEP) = 4.9; *χ*^2^(ANTIREG) = 6.76 (10) In this way, we could show that the nanodomains were registered across the bilayer leaflets.

However, let’s stay with this particular case for a while longer. By calculating a ratio between *χ*^2^(REG) and *χ*^2^(INDEP) we get a parameter that resembles the above introduced parameter RES, allowing us to characterize the experimental resolution between registered and independent nanodomains. By calculating the ratio also for the remaining two cases, we can compare the results in a more robust way: *χ*^2^(REG)/*χ*^2^(INDEP) = 2.53; *χ*^2^(REG)/ *χ*^2^(ANTIREG) = 3.58; *χ*^2^(INDEP)/*χ*^2^(ANTIREG) = 1.42. This comparison shows that registered vs. independent or registered vs. anti-registered nanodomains can be resolved more safely than independent from anti-registered nanodomains. This experimental result thus fully supports the results of the simulations, which identified the resolution between independent versus anti-registered nanodomains as the worst (**Figure 3**). In other words, the analysis shows a general trend evident for all Cases I, II and III, namely that it is easiest to distinguish between registered nanodomains and the other situations or between asymmetrically distributed domains versus the alternative scenarios, and what remains as the most challenging to distinguish is independent from anti-registered nanodomains.

### Case II: Accumulation of D and A in the nondomain region (*K*_D_(D) < 1 and *K*_D_(A) < 1) yields improved resolution

Since fluorescent probes that would be localized exclusively in one of the regions do not practically exist (*K*_D_(D/A) ≪ 1), we constrain our discussion here to the ones having *K*_D_(D, A) ∈ 〈0.1; 1〉. At the same time, we show more results for *K*_D_(D, A) ∈ 〈0.01; 1〉 in SI. For an efficient comparison of Cases I, II and III, it should also be noted that D and A having *K*_D_(D/A) = *X* or *K*_D_(D/A) = 1/*X*, where *X*is an arbitrary number, show the same affinity to the nanodomains or the regions outside of them.

Case II provides somewhat better resolution between registered and independent nanodomains in comparison to the remaining five situations. And as in the Case I, it appears experimentally the most challenging to resolve anti-registered from independent nanodomains and in this case also anti-registered nanodomains from nanodomains localized in one leaflet only (**Figure 5**). Generally, Case II appears more advantageous for the characterization of nanodomain coupling than Case I (compare **Figure 3** and **Figure 5** using the same value of *X*). The RES maps exhibit a strong ‘capital lambda’ shape indicating that the lower resolution is achieved for the nanodomains with *R*/*R*_0_ < 3.4 and Area < 15% (**Figure 5**). Although not so clearly, this characteristic shape is also evident for Case I (**Figure 3**). Overall, satisfactory resolution is reached for all the situations as early as for *K*_D_(D/A) = 0.2, thereby significantly expanding the selection of suitable D and A that can be used for the analysis (see (15) or **Table 2** for the most up-to-date list of suitable fluorescent probes).

### Example II: Bodipy-HDPC/fast DiI D/A pair (*K*_D_(D/A) = 0.1, *R*_0_ = 65 Å)

This D/A pair consiting of 1-hexadecanoyl-sn-glycero-3-phosphocholine labelled by Bodipy (donor) and DiI acceptor (**Table 3** and **Figure 4**) is characterized by the highest reported affinity for the nondomain region (**Table 2**) and large *R*_0_, therefore guarantiing the best resolution that can be reached nowadays when using ‘nondomain probes’. The pair has been used to characterize phase behaviour of giant unilamellar vesicles (22, 25) and belongs to popular D/A pairs in cell biology (26). Overall, this D/A pair exhibits good resolution characteristics that are comparable to the Bodipy-FL-GM1/Bodipy-564/570-GM1 D/A pair. More specifically, this and similar D/A pairs are suitable to resolve registered or asymterically distributed nanodomains from the alternative situations. Regarding the last ‘problematic’ scenario where independent nanodomains are being resolved from anti-registered ones, nanodomain coupling is under the detection limit only if nanodomains occupy less than 20 % of the membrane. Additionally, it becomes experimentally challenging to resolve anti-registered versus asymmetrically distributed nanodomains if they are smaller than 40 nm and occupy less than 10 % of the membrane (**Figure 4**).

### Case III: Accumulation of D and A in distinct bilayer regions (*K*_D_(D) > 1 and *K*_D_(A) < 1) yields the most robust resolution

In this case, high RES values are characteristic for all investigated situations (**Figure 6**) when using probes with a reasonably high affinity to the (non)domain region (*K*_D_(D) > 5 and *K*_D_(A) < 0.2). Specifically, RES reaches significantly higher values than for Case I and Case II (using the same value of *X* for this comparison) and remains high enough even for small nanodomains covering low fraction of the membrane (compare **Figure 3** and **Figure 5** with **Figure 6**). Although it persists as the most difficult to distinguish independent form anti-registered or anti-registered nanodomains from those localized asymmetrically in one leaflet as in Cases I and II, the resolution is perfectly adequate. Overall, considering that Case III provides good resolution for all the situations at *K*_D_(D) > 5 and *K*_D_(A) < 0.2, it appears as the most robust Case that should be used to guarantee stable high resolution for all scenarios that can arise (**Figure 1** A-B-C-D).

### Example III: NBD-DPPE/ Rhodamine-DOPE (*K*_D_(D) = 4.3 and *K*_D_(A) = 0.37, *R*_0_ = 64.1 Å)

This popular NBD-PE/Rh-PE FRET pair has been predominantly used in a vast majority of lipid mixing experiments in membrane fusion studies.(27, 28) Importantly, DPPE and DOPE lipids conjugated with NBD and Rhodamine, respectively, have also been used in a pioneering study by de Almeida et al, in which the authors discovered lipid nanodomains in artificial membranes.(21) Although the study did not give any accurate estimate regarding the nanodomain size, relative surface area occupied by nanodomains or inter-leaflet organization of nanodomains, the study represents, to our best knowledge, the first experimental study where FRET was used to detect membrane nanodomains.

As shown on **Figure 4**, this D/A pair would show excellent results in the studies of nanodomain organization, even though the affinity of this pair to the (non)domain region does not reach the maximum values that have been reported so far (compare the values on **Table 2**). More specifically, all coupling scenarios can be resolved from each other, with the exception of less likely situations when distinguishing independent vs. anti-registered nanodomains if Area < 10% or when distinguishing anti-registered vs. asymmetrically distributed nanodomains if *R* < 9 nm.

### Switching off *intra*-FRET. Does it lead to improved resolution?

In lipid bilayers, FRET occurs not only within one bilayer leaflet (*intra*-FRET) but also from one leaflet to the other one (*inter*-FRET).(10, 18, 29) Both processes take place simultaneously and independently of each other (**Figure 1**). However, since only *inter*-FRET can uncover how nanodomains are organized between opposite bilayer leaflets, we were interested whether the resolution could not be further improved by eliminating the possibility of *intra*-FRET. In the simulations, we therefore decided to switch off *intra*-FRET by placing D in one layer and A in the other layer and to investigate whether this will lead to improved resolution of MC-FRET. Thanks to recent experimental advances, a setup where D and A are located in opposite layers represents not only a hypothetical scenario but also a real experimental possibility. As shown in Doktorova et al,(30) asymmetric lipid bilayers, *i.e.* bilayers consisting of leaflets with different lipid composition, can be prepared by inter-vesicular exchange of outer leaflet lipids catalyzed by methyl-β-cyclodextrin. In this way, D in the outer leaflet could be replaced by A contained in exchange vesicles.

For better comparison, the results of simulations are presented in **Figure 7** where Cases I, II and III are represented by *K*_D_(D/A) = 5; *K*_D_(D/A) = 0.2; *K*_D_(D) = 5 and *K*_D_(A) = 0.2 respectively. Indeed, the resolution improves, but only for Case I where a satisfactory resolution is reached for *K*_D_(D/A) as low as 2, and only slightly for Case II. In contrast, the resolution remains constant or even gets a little worse for Case III. The degree of improvement is primarily determined by the extent to which *intra*-FRET is competitive for *inter*-FRET: the improvement is the most prominent for Case I where *intra*-FRET is extraordinary efficient due to entrapment of D and A within the nanodomains, and is insignificant for Case III, where D and A are separated from each other even when they are distributed symmetrically in both leaflets. Thus, by locating D and A into opposite leaflets, only the resolution for Case I is improved, and reaches similar results as the most robust Case III with the probes equally distributed between the leaflets.

### D/A Pairs With a Short Förster Radius Do Not Provide Sufficient Resolution

Finally, since it is widely known that the efficiency of FRET decreases quickly with the distance between a D and an A we set out to investigate the resolution of FRET as a function of the inter-layer distance *d.* This distance, defined as the distance at which D and A in one leaflet are transversally separated from D and A in the other leaflet (**Figure 8**), controls the efficiency of *inter*-FRET that is responsible for the sensitivity of FRET to the organization of nanodomains.

We simulated FRET for four different inter-layer distances: *d* ∈ {*R*_0_; *R*_0_/1.5; *R*_0_/2; *R*_0_/ 20}. By assuming a typical value of *R*_0_ = 58.7Å (see **Table 3**), D and A would be A) fully exposed to the bulk (*d* = *R*_0_); B) localized close to the lipid-water interface (*d* = *R*_0_/1.5); C) localized below the lipid-water interface (*d* = *R*_0_/2) or D) deeply buried in the membrane close to the bilayer centre (*d* = *R*_0_/20) (**Figure 8**). In this case, D and A basically occupy the same plane that is located at the bilayer centre (**panel D of Figure 8)**.

The analysis of **Figure 8** shows that the best resolution is obtained when D and A are located in the bilayer center. Although the resolution is declining slowly in the range *d* ∈ [*R*_0_/20; *R*_0_], any D/A pair whose *R*_0_ exceeds the bilayer thickness is perfectly suitable for the characterization of nanodomain coupling. As shown in the following example, since the resolution deteriorates rapidly for *d* > *R*_0_, care should be taken when selecting probes with a short *R*_0_.

### Example IV: DHE/Bodipy-PC(16:0) (*K*_D_(D) = 3.7 and *K*_D_(A) = 0.16, *R*_0_ = 28 Å)

This D/A pair exhibits a reasonable affinity for the (non)domain region, and thus represents a good D/A pair for the detection of nanodomains and determination of their size (13). At the same time, however, this pair has a short Förster radius, which reduces its sensitivity to the inter-leaflet organization of nanodomains (**Figure 4**). The final resolution depends on the vertical positions of DHE and Bodipy chromophores. Whereas the chromophore of Bodipy-PC is located close to the lipid-water interface (like the chromophores selected for Examples I-III)(16), the location of DHE is less clear. Nevertheless, under the extreme assumptions that DHE is located either at the lipid/water interface or close to the bilayer centre, the resolution appears poor in both cases (only the case when DHE is located at the interface is shown on **Figure 4**). This D/A pair is thus suitable only for the detection of nanodomains but less suitable in the research of nanodomain coupling.

### Design of an MC-FRET experiment and the applicability of the approach

Knowing the possibilities and limits of FRET, the next logical step is to use MC-FRET to determine the size, concentration, and inter-leaflet organization of nanodomains in real lipid membranes. Up to know, the method has been used in the membranes of giant unilamellar vesicles (GUVs)(8, 17, 23, 31, 32), as well as large unilamellar vesicles (LUVs)(31) and recently also in the membranes of giant plasma membrane vesicles (GPMVs)(7). In this last specific case, however, the study did not focus on the characterization of membrane nanodomains, but the quantification of protein dimerization in the membrane using the same MC-FRET methodology. Overall, success depends on determining as many parameters as possible that enter the simulation in an independent way. For clarity, we have summarized the most important parameters in **Table 4**, and as following texts implies, most of the parameters can be determined independently before starting the optimization procedure. The parameters including the Förster radius, the decay of donors in the absence of acceptors, the width of the lipid bilayer, but also the transversal localization of donor and acceptor chromophores within the lipid bilayer (16) (**Table 4**) can be determined/measured straightforwardly and input into the simulation prior to its start. In our research, we have used Bodipy-based chromophores, which are located universally close to the lipid/water interface (8, 16, 17). Thus, the parameters that remain to be determined comprise: the surface concentration of donors, *c*(D), and acceptors, *c*(A), in the membrane, the *K*_D_(D) and *K*_D_(A), the nanodomain radius 〈*R*〉, the surface density of nanodomains 〈*Area*〉 and, finally, the inter-leaflet organization of nanodomains.

**Table 4:**
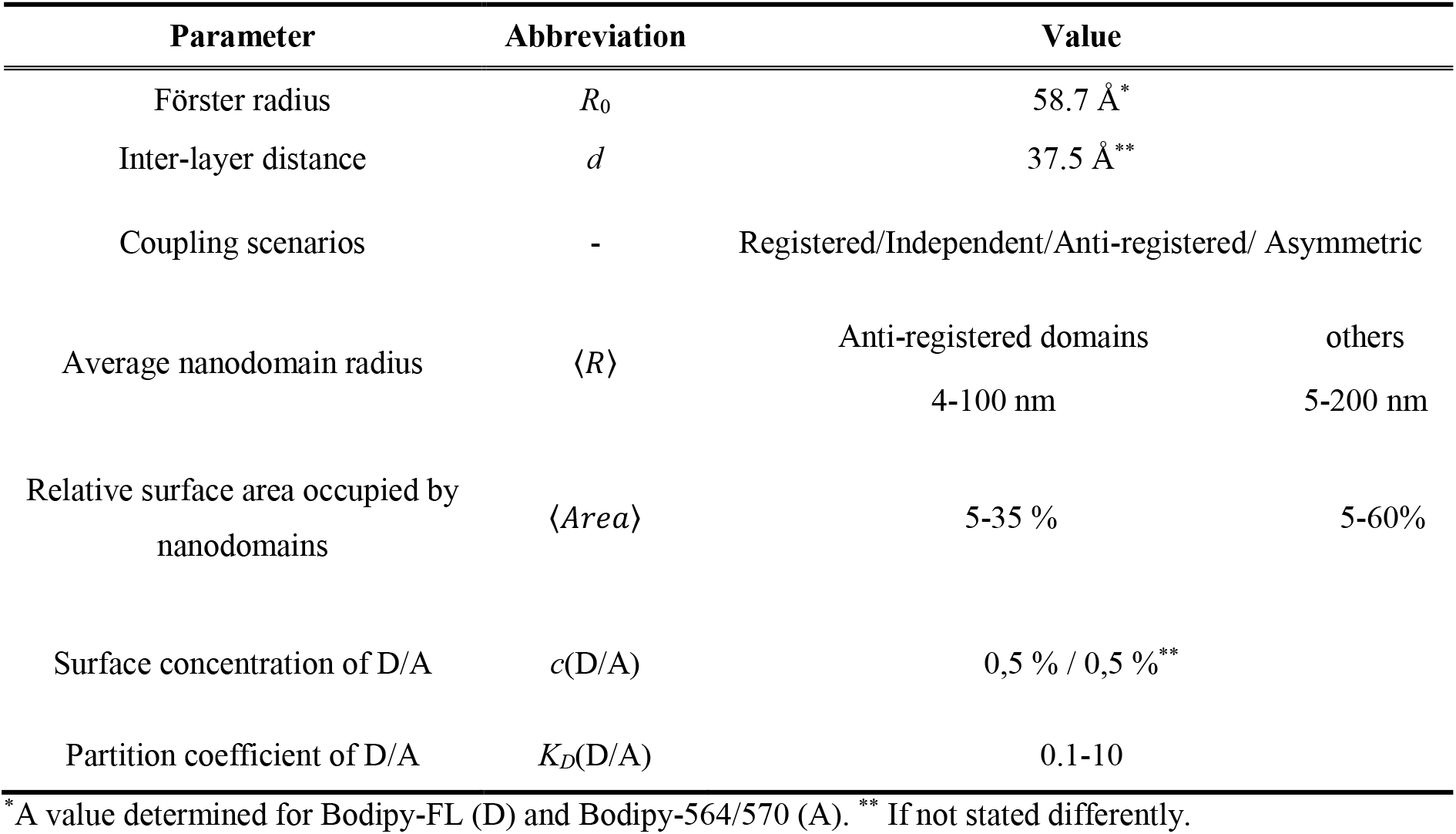
Input parameters entering the MC-FRET simulation

| Parameter | Abbreviation | Value |  |
| --- | --- | --- | --- |
| Förster radius | $R_0$ | 58.7 Å * | |
| Inter-layer distance | $d$ | 37.5 Å ** | |
| Coupling scenarios | - | Registered/Independent/Anti-registered/ Asymmetric |  |
| Average nanodomain radius | $\langle R \rangle$ | Anti-registered domains | others |
|  |  | 4-100 nm | 5-200 nm |
| Relative surface area occupied by nanodomains | $\langle Area \rangle$ | 5-35 % | 5-60% |
| Surface concentration of D/A | $c(D/A)$ | 0,5 % / 0,5 % ** | |
| Partition coefficient of D/A | $K_D(D/A)$ | 0.1-10 | |

\* A value determined for Bodipy-FL (D) and Bodipy-564/570 (A). \*\* If not stated differently.

As shown recently by Škerle et al (7), the concentrations *c*(D) and *c*(A) in the membranes of GPMVs or GUVs can be determined, for instance, by z-scan FCS by constructing a calibration curve that relates the intensity of D and A located in the membrane with the corresponding concentration. Alternatively, we developed an approach applicable to LUVs, GUVs and GPMVs that is based on the determination of *c*(D) and *c(A)* by FRET (15, 33). This approach builds upon the so-called Bauman-Fayer model, which provides the surface concentration (34) as one of the fitting parameters for homogeneous vesicles that have a composition as close as possible to that of nanoscopically heterogeneous membranes.

The robustness of the method will further increase if also *K*_D_(D) and *K*_D_(A) can be determined independently for the particular mixture of interest. This is, however, not always possible, but, very often, an MC-FRET experiment can be designed in such a way that the range of possible *K*_D_values is noticeably limited. E.g. when studying ganglioside nanodomains or nanodomains enriched by sphingomyelin, headgroup labeled gangliosides with exclusive localization in lipid domains (*K*_D_(D/A) ≥ 10) have proved useful. From the analyses carried out at that time, it emerged that the position of the global minima did not change over a wide range of *K*_D_(D/A) ≥ 5. In a similar study, we determined the size and concentration of cholera toxin nanodomains using Alexa-488-labeled Cholera toxin (donor, as expected, localized in the domains) and DiD acceptors excluded from those nanodomains with significantly ordered character (31). In an optimistic real-world scenario, the number of optimized parameters can thus be reduced to only 3: the nanodomain radius 〈*R*〉, the surface density of nanodomains 〈*Area*〉 and the inter-leaflet organization of nanodomains. In our fitting routine, we first scan the chi-squared space by sequentially changing 〈*R*〉 and 〈*Area*〉 for a given organization of nanodomains and then repeat the same procedure for the alternative coupling scenarios. In this way, we obtain a three-dimensional matrix of chi-squared values, in which we search for global and possibly other secondary minima. This procedure guarantees that we do not overlook similarly deep minima. For instance, in the lipid mixture consisting of DOPC/SM (90/10), we found two equally deep minima centered at 〈*R*〉 = 8 nm and 〈*Area*〉 = 37% and at 〈*R*〉 = 12 nm and 〈*Area*〉 = 55% (8) without knowing which of these minima corresponded to reality. In a less optimistic situation when the final *K*_D_ values are not known at all, it is still possible to repeat the entire fitting procedure for a set of different *K*_D_’s and try localize the global minima corresponding to the real values of the unknown parameters.

Purely out of our curiosity, we also carried out a completely different analysis as part of this work, described in detail in the SI. Briefly, we generated time-resolved fluorescence decays corresponding to the settings specified in **Table S1**. In the rest of the analysis, we treated these decays as experimentally recorded decays. We therefore fitted these decays using models considering registered, independent, anti-registered or asymmetrically distributed nanodomains (scenarios 1-4). In the final step, we compared the obtained chi-square values characterizing the quality of the fit as well as the input and output parameters of the simulation: 〈*R*〉, 〈*Area*〉 and possibly *K*_D_(D/A). This analysis ultimately shows that FRET does have the potential to characterize nanodomain coupling, as concluded in the paper. However, caution is warranted if there is no information on possible *K*_D_(D/A) values. In such a case, it cannot be ruled out that the analysis will provide several global minima as the final output, and it is then only up to the experimentalist to further narrow down the set of possible local minima.

The situation is noticeably more complicated in plasma membranes of living cells, in which the number of unknown parameters cannot be constrained as effectively as in the membranes of lipid vesicles. The local concentrations of D and A may vary, for instance due to local membrane curvature or local inhomogeneities, as well as it is generally more difficult to make any qualified estimate about the values of *K_D_*’s. Consequently, one may be less lucky in obtaining a reliable information about the coupling of nanodomains. But it is true that this approach will first need to be tested experimentally to find its limits when applied to cellular membranes. Of note, a reasonable compromise on the way from model to cellular membranes may consist of GPMVs. These vesicles have a flat membrane and as shown in Skerle et al (7), the list of unknown parameters that enter the simulation may be shortened considerably.

## CONCLUSIONS: CRITICAL PARAMETERS THAT CONTROL THE RESOLUTION & SUITABLE D/A PAIRS

The analysis presented in this article identified the following experimental parameters as the most critical in the characterization of nanodomain coupling:

First, the Förster radius and its value related to the inter-layer distance *d,* at which D and A in one leaflet are transversally separated from D and A in the other leaflet (**Figure 8**). Depending on this parameter, the resolution appears relatively stable in the range *d* ∈ [*R*_0_/20; *R*_0_/1.5]: it is the best when D and A are located in the bilayer center, *i.e.* if *d* ≪ *R*_0_ (see **Figure 8**) and declines sharply beyond this range. Since most of fluorescent probes used nowadays are located at the lipid-water interface, *R*_0_ should not be lower than the bilayer thickness.

Second, the resolution of FRET is controlled by the affinity of D and A to the (non)domain region. Although the affinity of available probes is generally low (**Table 2**), the *K*_D_’s are high/low enough to allow for characterization of nanodomain organization with unprecedented detail. More specifically, D and A having moderate affinity to the nanodomains (*K*_D_(D/A) > 5) or the nondomain region (*K*_D_(D/A) < 0.2) can resolve registered from independent, anti-registered or asymmetrically organized nanodomains as well as asymmetrically distributed nanodomains from independent or anti-registered ones in a broad range of nanodomain radii *R*/*R*_0_ ≥ 2 and relative surface area occupied by the nanodomains *Area* ≥ 10% (compare **Figure 3**, **Figure 5** and **Figure 6**). At the same time, it appears experimentally challenging to distinguish independent from anti-registered nanodomains, since, in this case, nanodomains are distributed over the bilayer surface in a very similar manner. Of all three Cases I, II and III analyzed, Case III where D and A exhibit opposite reasonably high affinity to the nanodomains and the region outside of them (*K*_D_(D) > 5 and *K*_D_(A) < 0.2) provides the best resolution.

Third, the resolution of the method depends on the extent to which *inter*-FRET is competitive with *intra*-FRET and, as expected, the method performs worst if *intra*-FRET is dominant. This is most evident in Case I (*K*_D_(D) > 1 and *K*_D_(A) > 1) where D and A are located in the nanodomains. In this case, however, the resolution can be improved by placing D and A into opposite bilayer leaflets, thereby eliminating *intra*-FRET.

According to our analysis, the best performance offer Bodipy-FL-GM1/Bodipy-564/570-GM1 (*K*_D_(D/A) ≥ 20, *R*_0_ = 58.4Å) or NBD-DPPE/ Rhodamine-DOPE (*K*_D_(D) = 4.3 and *K*_D_(A) = 0.37, *R*_0_ = 64.1 Å) D/A pairs. Both D/A pairs reach a satisfactory resolution, except of extreme conditions when nanodomains are small (nanodomain radius < 10 nm or *R*/*R*_0_ = 3.4) or occupy a small part of the lipid bilayer (surface density of nanodomains < 10%). Overall, MC-FRET appears as a robust method that - when using D/A pairs with good characteristics - yields otherwise difficult-to-reach characteristics of membrane lipid nanodomains.

## AUTHOR CONTRIBUTIONS

RS conceived the idea, BC and RS developed the simulation code, BC performed the simulations and analysed the data with help of DD. BC and RS interpreted the data and wrote the manuscript. All authors revised the manuscript.

## ACKNOWLEDGEMENTS

RŠ and BC and DD acknowledge GAČR Grant 20-01401J. RŠ acknowledges European Union’s Horizon 2020 research and innovation program under Grant Agreement No. 101017902. BC would like to acknowledge Project No. SVV 260586 of the Charles University. Computational resources were supplied by the project “e-Infrastruktura CZ” (e-INFRA CZ LM2018140) supported by the Ministry of Education, Youth and Sports of the Czech Republic. We thank prof. Martin Hof for reading the manuscript and providing useful comments.

