## Supplemental Information for "Inter-leaflet Organization of Membrane Nanodomains: What Can(not) Be Resolved by FRET?"

### RESULTS: ANALYSING THE RESOLUTION OF FRET

**Figures S1** is an extension of **Figure 4** in the main text. It shows the resolution diagrams for the probes with high affinity to nanodomains ($K_{\mathrm{D}}(\mathrm{D,A}) \in \langle 10; 1000\rangle$). It is evident from the resolution maps that the resolution improves only slightly in the range $K_{\mathrm{D}}(\mathrm{D,A}) \in \langle 10; 100\rangle$ and is constant beyond $\mathrm{K_D}(\mathrm{D/A}) > 100$.

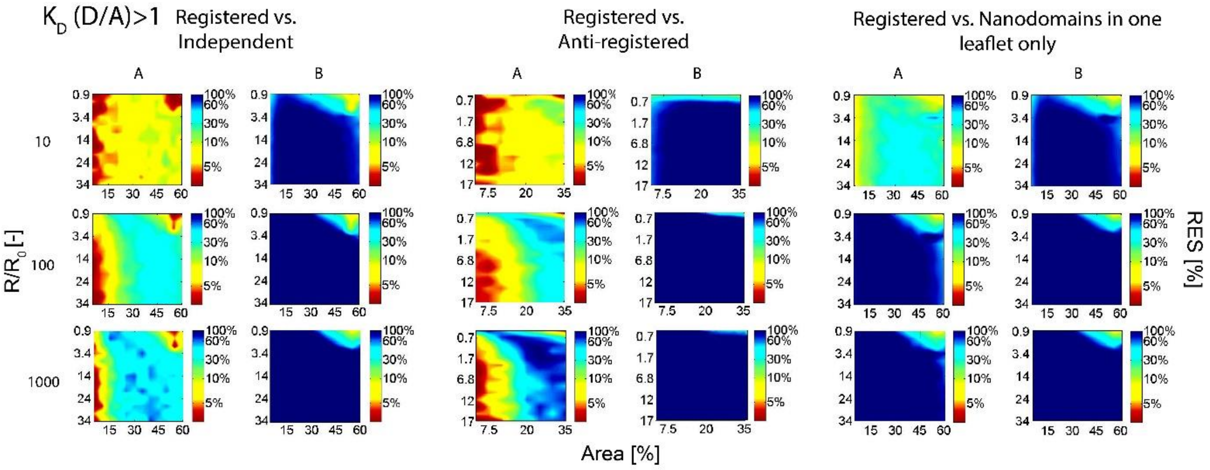


**Figure S1:** RES diagrams for D and A exhibiting high affinity to the nanodomains ($K_{\mathrm{D}}(\mathrm{D,A}) \in \langle 10; 1000\rangle$). (A) D and A were located in both bilayer leaflets or (B) they were located in the opposite leaflets.

**Figures S2** is an extension of **Figure 6** in the main text. It shows the resolution diagrams for the probes with high affinity to the nondomain phase ($K_{\mathrm{D}}(\mathrm{D,A}) \in \langle 0.001; 0.1\rangle$). It is evident from the resolution maps that the resolution improves only slightly in the range $K_{\mathrm{D}}(\mathrm{D,A}) \in \langle 0.01; 0.1\rangle$ and is constant if $\mathrm{K_D}(\mathrm{D/A}) < 0.01$.

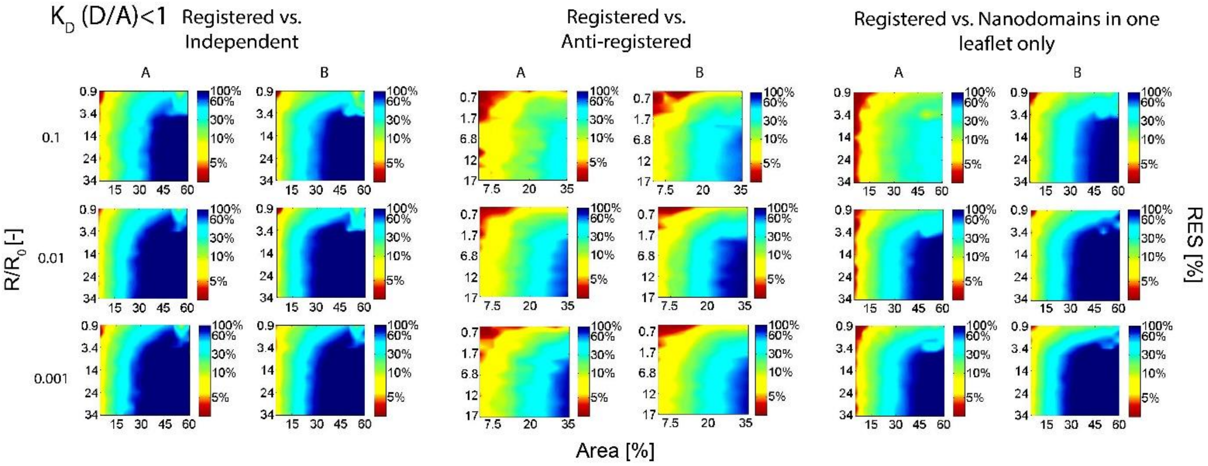


**Figure S2:** RES diagrams for D and A exhibiting high affinity to the nondomain phase ($K_\mathrm{D}(\mathrm{D},\mathrm{A}) \in \langle 0.001; 0.1 \rangle$). (A) D and A were located in both bilayer leaflets or (B) they were located in the opposite leaflets.

**Figure S3** shows the dependence of the FRET resolution towards nanodomain coupling shown for a set of different acceptor to lipid ratios. The dependence is relatively flat between the acceptor to lipid ratio 1:200 and 1:1000. Thus, any value selected within this range guarantees good FRET resolution.

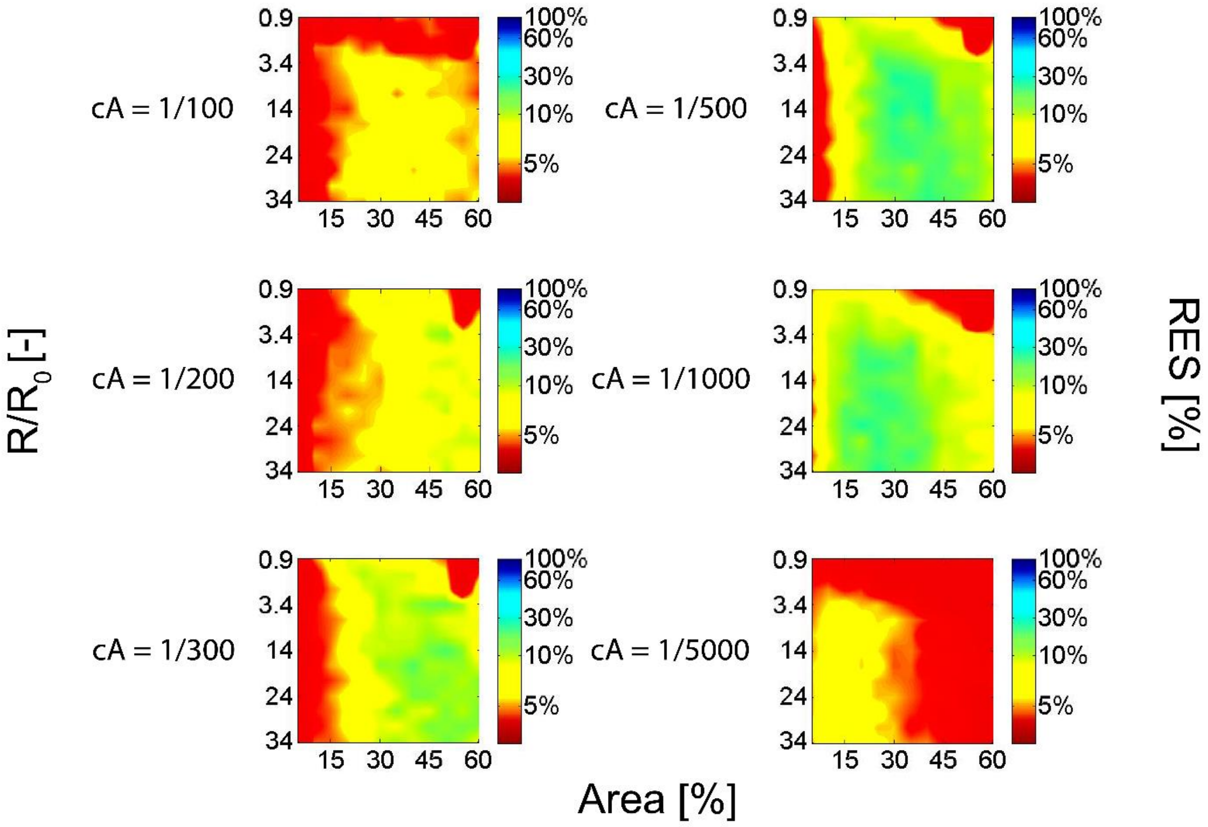


**Figure S3:** RES diagrams characterizing the resolution of FRET between registered and independent nanodomains shown for a set of different acceptor to lipid ratios. $K_D(D/A) = 10$.

As part of this study, we also performed the following test verifying the potential of MC-FRET to characterize nanodomain coupling. We generated additional time-resolved fluorescence decays corresponding to the settings specified in **Table S1**. In the rest of the analysis, we treated these decays as experimentally recorded decays. We therefore fitted these decays using models considering registered, independent, anti-registered or asymmetrically distributed nanodomains (scenarios 1-4). In the final step, we compared the obtained chi-square values characterizing the quality of the fit as well as the input and output parameters of the simulation: *R*, *Area* and possibly $K_D$.

Altogether, we have performed three tests, each of them mimicking a different situation: In the first test, we assumed that prior to the start of the FRET experiment, the Kd values are known. In the test, we generated a total of three curves declared as experimental: a decay corresponding to D and A undergoing FRET in the presence of A) registered nanodomains ($R$ = 60 nm, *Area* = 50 %, $K_D$ = 5); or B) independent nanodomains ($R$ = 60 nm, *Area* = 50 %, $K_D$ = 0.1). According to the RES diagram shown on **Figures 4 and 5**, FRET should achieve reasonable resolution under such set conditions. And C) FRET taking place in the bilayer with anti-registered nanodomains ($R$ = 30 nm, *Area* = 25 %, $K_D$(D) = 10 and $K_D$(A) = 0.1). In this case, the conditions have been chosen to correspond to the situation when FRET has a limited capacity to characterize nanodomain coupling.

Since the $K_D$ values are a priori known in Test 1, we optimized only $R$ and *Area* and then compared the fits as well as the input and output parameters. Regarding the analysis based on the 'experimental' decay A), the 'experimental' decay B) and the 'experimental' decay C), it can be seen from **Table S2** that the lowest chi-square value of all four possible scenarios, was obtained for registered (decay A), independent (decay B) or anti-registered (decay C) nanodomains. Importantly, the input and output $R$ and *Area* values match perfectly. It is perhaps worth mentioning the relatively shallow chi-square minimum obtained for decay A), which substantially increases the uncertainty in determining the resulting nanodomain size and the relative area occupied by nanodomains (**Table S2**).

The second test was intended to imitate a situation where, before starting the experiment, it is known whether the used donors and acceptors have a reduced or increased affinity to the nanodomains, without knowing the specific value. Thus, we took the 'experimental' decays A) and C) and fitted it back by the models assuming registered-independent-anti-registered or asymmetrically distributed nanodomains and the following $K_D$ values 2-5-10 and combinations of $K_D$(D,A) 10-0,1; 5-0,2; 2-0,5. As can be seen from **Table S3**, even in this case the position of the global minimum coincides perfectly with the input data ($R$ = 60 nm, *Area* = 50 %, $K_D$ = 5 in case of registered experimental conditions and $R$ = 30 nm, *Area* = 25 %, $K_D$(D) = 10 and $K_D$(A) = 0.1 for the unregistered settings).

Finally, the third test was conducted under the assumption that no a priori knowledge about the $K_D$ values is available. For simplicity, we mimicked this situation by repeating the procedure described in Test 2. However, in this final test, we have substantially expanded the range of possible $K_D$ values summarized in **Table S4**. The selected $K_D$ values try to cover all possibilities: increased affinity of both D and A to nanodomains, increased affinity of both D and A to the nondomain region and the situation when D and A have opposite affinity the domain and nondomain regions. Even in this test, despite a significantly larger number of possibilities, where different combinations of optimized parameters can theoretically lead to an equally good fit, the performed analysis provides one global minimum.

Overall, this analysis ultimately shows that FRET indeed has the potential to characterize nanodomain coupling, as concluded in the paper. However, caution is warranted if there is no information on possible $K_{\mathrm{D}}(\mathrm{D/A})$ values. In such a case, it cannot be ruled out that the analysis will provide several global minima as the final output, and it is then only up to the experimentalist to further narrow down the set of possible local minima.

**Table S1:** Input parameters entering the simulation. This simulation generates time-resolved fluorescence decays that are treated in the analysis as experimentally recorded decays.

| exp. decay | Settings | $K_D$(D,A) | R (nm) | Area |
|---|---|---|---|---|
| A | Registered | 5 | 60 | 50% |
| B | Antiregistered | 10-0.1 | 30 | 25% |
| C | Independent | 0.1 | 60 | 50% |

**Table S2:** Simulation output parameters received as part of Test 1

| exp. Decay A - REGISTERED | | | | exp. decay B - ANTIREGISTERED | | | |
|---|---|---|---|---|---|---|---|
| model | chi-squared | R (nm) | Area | model | chi-squared | R (nm) | Area |
| REG | 1.025 | 45±15/145±25 | 50±5% | REG | 1.140 | 8±3 | 6±3% |
| ANT | 3.126 | 30±10 | 43±7% | ANT | 1.040 | 45±15 | 20±5% |
| IND | 2.139 | 21±5 | 55±5% | IND | 1.106 | 8±1 | 23±3% |
| ASY | 6.580 | 40±20 | 43±7% | ASY | 1.076 | 20±10 | 45±5% |

| exp. decay C - INDEPENDENT | | | |
|---|---|---|---|
| model | chi-squared | R (nm) | Area |
| REG | 6.157 | 18±2/120±40 | 35±15% |
| ANT | 116.317 | 21±5 | 27±3% |
| IND | 1.270 | 55±25 | 53±7% |
| ASY | 10.352 | 23±7 | 53±7% |

**Table S3:** Simulation output parameters received as part of Test 2

| experimental decay A - REGISTERED | | | | | | | |
|---|---|---|---|---|---|---|---|
| fitting model - REGISTERED | | | | fitting model - ANTIREGISTERED | | | |
| $K_D$(D,A) | chi-squared | R (nm) | Area | $K_D$(D,A) | chi-squared | R (nm) | Area |
| 10 | 1.082 | 21±1 | 57±3% | 10 | 1.252 | 30±10 | 30±5% |
| 5 | 1.025 | 45±15/145±25 | 50±5% | 5 | 3.126 | 30±10 | 43±7% |
| 2 | 7.352 | 53±27 | 55±5% | 2 | 8.634 | 30±11 | 57±3% |
| fitting model - INDEPENDENT | | | | fitting model - ASYMMETRIC | | | |
| $K_D$(D,A) | chi-squared | R (nm) | Area | $K_D$(D,A) | chi-squared | R (nm) | Area |
| 10 | 1.153 | 25±10 | 35±5% | 10 | 4.242 | 25±5 | 40% |
| 5 | 2.139 | 21±5 | 55±5% | 5 | 6.580 | 40±20 | 43±7% |
| 2 | 11.492 | 7±3 | 57±3% | 2 | 9.234 | 30±5 | 43±7% |
| experimental decay B - ANTIREGISTERED | | | | | | | |
| fitting model - REGISTERED | | | | fitting model - ANTIREGISTERED | | | |
| $K_D$(D,A) | chi-squared | R (nm) | Area | $K_D$(D,A) | chi-squared | R (nm) | Area |
| 10-0.1 | 1.140 | 8±3 | 6±3% | 10-0.1 | 1.040 | 45±15 | 20±5% |
| 5-0.2 | 1.103 | 7±1 | 9±1% | 5-0.2 | 1.151 | 65±35 | 16±4% |
| 2-0.5 | 1.482 | 140±40 | 45±5% | 2-0.5 | 10.447 | 22±2 | 20±5% |
| fitting model - INDEPENDENT | | | | fitting model - ASYMMETRIC | | | |
| $K_D$(D,A) | chi-squared | R (nm) | Area | $K_D$(D,A) | chi-squared | R (nm) | Area |
| 10-0.1 | 1.106 | 8±1 | 23±3% | 10-0.1 | 1.076 | 20±10 | 45±5% |
| 5-0.2 | 1.203 | 12±3 | 33±17% | 5-0.2 | 1.221 | 115±35 | 50±10% |
| 2-0.5 | 1.627 | 20±5 | 43±2% | 2-0.5 | 2.637 | 20±10 | 50±10% |

**Table S4:** Simulation output parameters received as part of Test 3.

| experimental decay A - REGISTERED domains | | | | | | | |
|---|---|---|---|---|---|---|---|
| fitting model - REGISTERED | | | | fitting model - ANTIREGISTERED | | | |
| $K_D$(D,A) | chi-squared | R (nm) | Area | $K_D$(D,A) | chi-squared | R (nm) | Area |
| 10 | 1.082 | 21±1 | 57±3% | 10 | 1.252 | 30±10 | 30±5% |
| 5 | 1.025 | 45±15/145±25 | 50±5% | 5 | 2.988 | 30±10 | 43±7% |
| 2 | 7.352 | 53±27 | 55±5% | 2 | 8.634 | 30±11 | 57±3% |
| 0.1 | 1.113 | 6±1 | 57±3% | 0.1 | 8.881 | 2±1 | 45±10% |
| 0.2 | 1.046 | 53±27 | 55±5% | 0.2 | 9.124 | 2±1 | 40±10% |
| 0.5 | 6.193 | 30±5 | 53±7% | 0.5 | 3.126 | 30±10 | 43±7% |
| 10-0.1 | 30.979 | 5±1 | 53±3% | 10-0.1 | 9.641 | 3±2 | 40±10% |
| 5-0.2 | 25.021 | 5±1 | 50±5% | 5-0.2 | 9.623 | 3±2 | 37±7% |
| 2-0.5 | 12.802 | 6±2 | 6±2% | 2-0.5 | 9.663 | 35±5 | 45±15% |
| fitting model - INDEPENDENT | | | | fitting model - ASYMMETRIC | | | |
| $K_D$(D,A) | chi-squared | R (nm) | Area | $K_D$(D,A) | chi-squared | R (nm) | Area |
| 10 | 1.153 | 25±10 | 35±5% | 10 | 4.242 | 25±5 | 40% |
| 5 | 2.139 | 21±5 | 55±5% | 5 | 6.580 | 40±20 | 43±7% |
| 2 | 11.492 | 7±3 | 57±3% | 2 | 9.234 | 30±5 | 43±7% |
| 0.1 | 9.455 | 6±1 | 55±5% | 0.1 | 3.992 | 55±25 | 50±5% |
| 0.2 | 11.564 | 6±1 | 55±5% | 0.2 | 5.739 | 45±15 | 50±5% |
| 0.5 | 12.236 | 7±2 | 6±1% | 0.5 | 9.071 | 20±10 | 45-55% |
| 10-0.1 | 18.455 | 5±1 | 7±3% | 10-0.1 | 10.354 | 3±1 | 50±10% |
| 5-0.2 | 15.010 | 6±1 | 7±3% | 5-0.2 | 10.640 | 3±1 | 50±10% |
| 2-0.5 | 13.167 | 6±1 | 7±3% | 2-0.5 | 10.118 | 2±1 | 30±10% |
